# Interaction between theta-phase and spike-timing dependent plasticity simulates theta induced memory effects

**DOI:** 10.1101/2021.11.24.469900

**Authors:** Danying Wang, George Parish, Kimron L. Shapiro, Simon Hanslmayr

## Abstract

Rodent studies suggest that spike timing relative to hippocampal theta activity determines whether potentiation or depression of synapses arise. Such changes also depend on spike timing between pre- and post-synaptic neurons, known as spike-timing-dependent plasticity (STDP). STDP, together with theta-phase-dependent learning, has inspired several computational models of learning and memory. However, evidence to elucidate how these mechanisms directly link to human episodic memory is lacking. In a computational model, we modulate long-term potentiation (LTP) and long-term depression (LTD) of STDP, by opposing phases of a simulated theta rhythm. We fit parameters to a hippocampal cell culture study in which LTP and LTD were observed to occur in opposing phases of a theta rhythm. Further, we modulated two inputs by cosine waves with synchronous and asynchronous phase offsets and replicate key findings in human episodic memory. Learning advantage was found for the synchronous condition, as compared to the asynchronous conditions, and was specific to theta modulated inputs. Importantly, simulations with and without each mechanism suggest that both STDP and theta-phase-dependent plasticity are necessary to replicate the findings. Together, the results indicate a role for circuit-level mechanisms, which bridges the gap between slice preparation studies and human memory.

**Author Summary:** Long-lasting changes in synaptic connectivity between neurons have been suggested to support learning and memory processes at the cellular level in the brain. Such synaptic modifications depend on synchronous activation of neurons, which leads to generate brain oscillations. Human memory studies focus on the relationships between brain oscillations and memory processes. Direct evidence on how the cellular mechanism links to human memory behaviour is lacking. To investigate the direct link between synaptic plasticity mechanisms and human memory formation, we built a computational neural network that implements two synaptic plasticity mechanisms, which are well-established in the rodents’ hippocampus. One mechanism shows that strengthening or weakening in synaptic connectivity depends on the phases of ongoing brain oscillation at theta frequency (4 – 8 Hz), which is a dominant signal in the hippocampus. The other mechanism suggests that synaptic modification depends on the precise timing of action potentials between two neurons. Our model successfully reproduces results from rodents, as well as several human episodic memory studies which demonstrated that human associative memory performance depends on phase synchronisation in theta frequency. These findings suggest a link between specific learning mechanisms at cellular level and human memory behaviour.

## Introduction

Modification of the strength of synaptic connections is suggested to be a key component of the mechanism underlying human memory. Such synaptic modification depends on synchronizing activity between neurons (Hebb, 2002; Markram et al., 1997). Therefore, brain oscillations that synchronise neuronal activity might be crucial for memory processes. Synchronized theta oscillatory activity (4 – 8 Hz), a dominant rhythm in the hippocampus, is thought to play a key role in synaptic modification (Buzsáki, 2002). Rodent studies show that timing of inputs relative to a theta cycle are important to long-term potentiation (LTP) and long-term depression (LTD) of hippocampal synapses. In vitro, LTP can be induced by a brief burst of pulses delivered at the CA1 theta peak while LTD occurs if the pulse is delivered at theta trough (Huerta & Lisman, 1995). Such theta-dependent synaptic plasticity has also been observed *in vivo* in CA1 (Hölscher et al., 1997; Hyman et al., 2003) as well as in the dentate gyrus (Pavlides et al., 1988).

Inspired by the precise timing that hippocampal theta creates for synaptic modification, several neural models have been proposed to investigate the role of hippocampal theta dynamics in learning and memory (Hasselmo et al., 2002; Ketz et al., 2013; Norman et al., 2006; Parish et al., 2018). One of these models implemented theta phase reversal between mono-synaptic and tri-synaptic pathways, i.e. from entorhinal cortex (EC) to CA1 via Schaffer collaterals, with the aim of promoting efficient learning by separating memory formation into phases of encoding and retrieval (Hasselmo et al., 2002; Ketz et al., 2013). The other two models purport to explore the theoretical role that theta might play in network plasticity and stability, by limiting the occurrence of LTP to a phase of high inhibition and the occurrence of LTD to a phase of low inhibition, allowing for an efficient encoding of new associations and the protection of pre-existing memories from interference(Norman et al., 2006; Parish et al., 2018). The present modelling work attempts to explain the relationship more fully between theta and synaptic modification at the neuronal level, based upon observations of theta dependent plasticity in hippocampal cells (Huerta & Lisman, 1995).

Based on physiological studies and computational neural modelling, synchronising inputs (i.e. arriving at the same time in the hippocampus) to the LTP inducing theta phase should lead to successful encoding of the association between those inputs, as compared to when inputs are desynchronised (i.e. arrive at different times in the hippocampus). Recently, this assumption has been demonstrated by human episodic memory studies. In these studies (Clouter et al., 2017; Wang et al., 2018), luminance of videos and amplitude of sounds were modulated at 4 Hz to entrain visual and auditory cortical activity at theta frequency. Participants were asked to recall the correctly paired video when they were cued with a sound. The recall accuracy for synchronously presented stimuli was significantly better than the recall accuracy for asynchronously presented stimuli. More importantly, this synchronisation induced memory advantage was only shown for the theta modulated stimuli; not for the faster (alpha, 10.5 Hz) or slower (delta, 1.7 Hz) frequency modulated stimuli. This memory effect goes beyond multisensory binding at purely perceptual levels, which is demonstrated by significantly better recall in the theta synchronous condition compared to a condition where stimuli were unmodulated (Clouter et al., 2017).

The human theta-induced memory effect provides evidence for the causal role of theta phase synchronisation in episodic memory formation, indicating a common mechanism of hippocampus-dependent memory behaviour between human and rodent memory models. Interestingly, both human studies support the idea that more effective memory formation depends on finely tuned timing between inputs. The recall accuracy in the asynchronous conditions was not modulated by the degree of asynchrony, that is, recall accuracy was similar across all phase offset conditions for 4 Hz (i.e. same performance levels for 90°, 270°, or 180°; Clouter et al., 2017; Wang et al., 2018). An alternative explanation would be that the net hippocampal theta power increases because of the entrained cortical inputs, which cannot explain why the performance in the 90° and 270° conditions did not get a benefit compared to the 180° condition. One such finely tuned timing mechanism in synaptic plasticity is spike timing dependent plasticity (STDP). This theoretical framework suggests that synaptic efficacy decays exponentially as a function of the delay between spikes of pre-synaptic and post-synaptic neurons, which leaves a very narrow time window (∼25 ms) for efficient synaptic weight changes (Song et al., 2000). The sign of synaptic modification, LTP or LTD depends on the temporal order of the two spikes. If the post-synaptic neuron fires in a short delay after the firing of the pre-synaptic neuron, then the synaptic connection from the pre-synaptic neuron to the post-synaptic neuron will be strengthened. LTD of the synapses will be induced if the order of spikes is reversed. Therefore, computational modelling of the STDP learning rule suggests that STDP is more biophysically feasible, as compared to Hebbian learning because it provides a solution to competitive synapses that share the same post-synaptic neurons (Song et al., 2000). Empirical evidence of STDP has also been shown in several slice and cell culture studies in rodent hippocampus (Bi & Poo, 2001), as well as in human hippocampus, although LTP was induced in a wider temporal window of pre- and post-synaptic spike timing, between -80 ms and +10 ms (Silva et al., 2010).

The aim of the present computational study is to better understand the role of the two components, theta-phase-dependent plasticity and STDP, in human episodic memory formation. To this end we build on a previous model, the Sync/deSync model (Parish et al., 2018) and create a new version of the model that enables the STDP learning rule to modify hippocampal weight changes. Significantly, we separate LTP and LTD processes, which are now modulated by opposing theta phases in the hippocampus, where previously plasticity was more generally modulated by theta phase (i.e. both LTP & LTD). Here, LTP is induced at an inhibitory phase of theta, and LTD at the excitatory phase. In the rodent, theta activity can be measured from fissure EEG. Synapses are strengthened via LTP when the input from EC is strongest at theta trough. Whereas at the theta peak of fissure EEG, synaptic input from EC is weak and neurons in the CA1/CA3 are more active, LTD occurs (Hasselmo, 2005). To find the best set of parameters, we first aimed to reproduce findings from Huerta and Lisman (1995) showing that LTP and LTD occur in opposing theta phases. We then used these parameters to reproduce findings from the human episodic memory studies of Clouter et al. (2017) and Wang et al. (2018). We aimed to simulate the important results that the theta phase synchrony induced memory effect is specific to theta modulated stimuli and goes beyond perceptual binding, which is indicated by better learning in the theta synchronous condition than in the unmodulated condition. Moreover, to demonstrate that both STDP and theta-phase-dependent plasticity are necessary to simulate the behavioural findings, we evaluated model performance by comparing the full model with two partial versions of the model that switch one component off and leave the other on. Our findings provide a computational framework to link human behaviour to hippocampal function at the circuitry or even cellular level.

## Methods

### Modelling principles and experimental paradigm

Inspired by models implementing reversal phasic relationships of the theta rhythm in hippocampal subfields (Hasselmo, 2005; Ketz et al., 2013), we build on a previous model, the Sync/deSync model (Parish et al., 2018). This previous model focuses on the key function of CA1, where theta rhythm establishes an inhibitory phase (i.e. suppressed neural firing in CA1) during which time synapses undergo LTP, and a facilitatory phase (i.e. enhanced neural firing in CA1) when LTD occurs.

The model parameters were fit in relation to a hippocampal cell culture study (Huerta & Lisman, 1995) showing that stimulation at opposing theta phases induces LTP and LTD, respectively. Further, we simulated human episodic memory experiments involving a multisensory entrainment paradigm (Clouter et al., 2017; Wang et al., 2018) that showed human episodic memory formation depends on stimulus input timing relative to theta oscillations. In these studies, the luminance of videos and amplitude of sounds were modulated by 4 Hz cosine waves. The phase offsets of modulated sounds were either in-phase (0°) or out-of-phase (90°, 180°, and 270°) from modulated videos (Figure 1B). Each pair of sounds and videos was presented for three seconds while participants were asked to form associations between them. To test participants’ memory on the associations between the videos and sounds, each sound was presented again during recall and participants were asked to select the video that was paired with the sound earlier in the study phase. To simulate this paradigm, we fed two 4 Hz cosine waves as visual and auditory stimulus inputs into two independent neural populations. The phase offsets of the auditory stimulus are either in-phase (0°) or out-of-phase (90°, 180°, and 270°) from the visual stimulus input.

**Figure 1.**
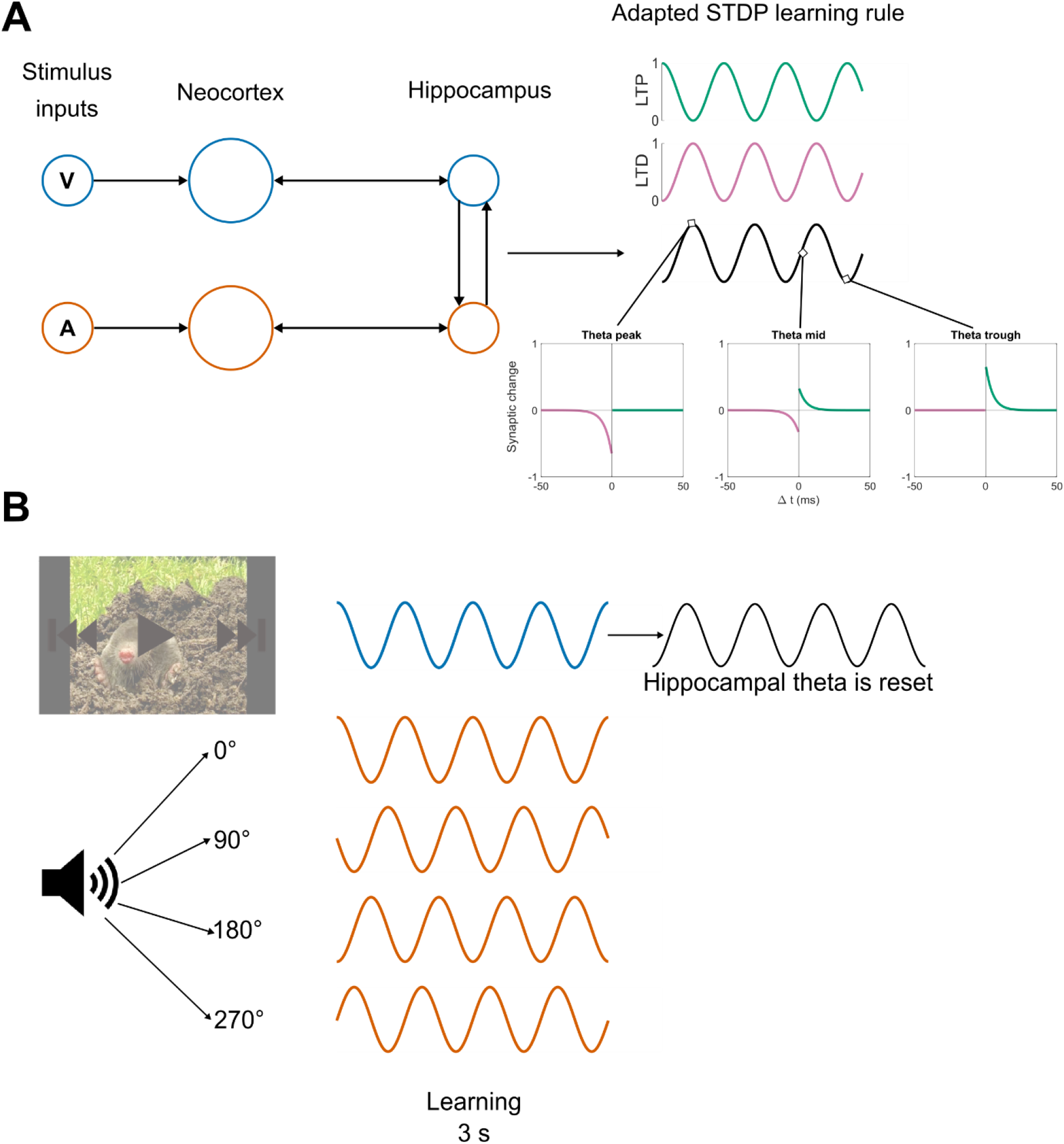
Model Architecture and Experimental Paradigm. ***A,*** Two subgroups of neurons are created to represent the visual and auditory stimulus, respectively, in both NC and the hippocampus. STDP is enabled and modulated by on-going theta oscillations, which modulates LTP (green) and LTD (purple) at opposing phases. ***B,*** to simulate the human episodic memory experiments using a multisensory entrainment paradigm (Clouter et al., 2017; Wang et al., 2018), two 4 Hz cosine waves as visual (blue) and auditory (orange) stimulus inputs were fed into two independent neural populations. The phase offsets of the auditory stimulus are either in-phase (0°) or out-of-phase (90°, 180°, and 270°) from the visual stimulus input. Hippocampal theta phase was reset with a 180° offset from modulated visual stimulus after stimulus onset.

### Neuron physiology

The model architecture is carried forward from that of the Sync/deSync model (Parish et al., 2018), which consists of two groups of neurons that represent the neo-cortex (NC) and hippocampus. Where previously these groups were themselves split into subgroups to denote different image-based stimuli (Parish et al., 2018), here two subgroups of neurons are created to represent the visual and auditory stimulus, respectively, in both NC and the hippocampus (Figure 1A). The total number of neurons in NC was 20 and the number of neurons in the hippocampus was 10.

Neuron membrane potential changes are modelled by an integrate-and-fire equation (Equation 1), where the membrane potential decays over time to a resting potential (E_L_ = - 70mV) at a rate dictated by the membrane conductance (g_m_ = 0.03). Here, a spike event is generated if the voltage exceeds a threshold (V_th_ = -55mV), at which time the voltage is clamped to the resting potential for an absolute length of time to approximate a refractory period (V_ref_ = 2ms). As well as the leak current, the input current for model neurons contains the sum of all spike events occurring at pre-synaptic neurons (I_syn_), alternating current (AC) that represents NC alpha and hippocampal theta oscillations (I_AC_), any existing direct current (I_DC_) and an after-depolarisation (ADP) function (I_ADP_), described later.

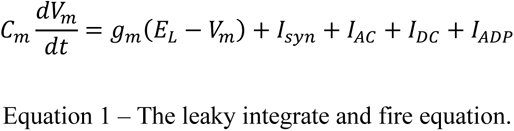

Equation 2 explains the process by which neurons communicate through spike events. Here, an excitatory post-synaptic potential (EPSP) provided an additive exponential function that diminishes the further the current time point (*t*) is form the initiating spike event (t_fire_). The amplitude of the function is dictated by the current potentiation of the post-synaptic synapse (0≤ *ρ* ≤1) multiplied by its maximal weight (W_max_). All spike events had a delay of 2ms before they reached post-synaptic connections.

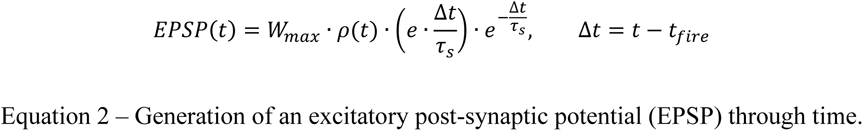

Hippocampal neurons received additional input from an ADP function, as in previous models (Jensen et al., 1996; Parish et al., 2018; Equation 3; A_ADP_ = 0.2nA, τ_ADP_ = 250ms). This provided exponentially ramping input, which was reset after each spike-event (t_fire_). The ADP function bestowed upon model hippocampal neurons an inherent preference for spiking at a theta frequency, as well as slowing down their overall firing rate as a source of effective inhibition. It was this latter function that played a central role in the previous iteration of the Sync/deSync model (Parish et al., 2018), where the desynchronisation of an oscillating cell assembly required more input for slower spiking neurons.

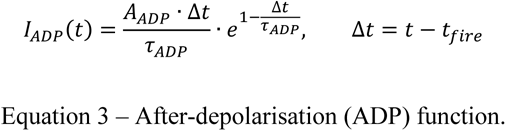

The learning rule was implemented via an adapted spike-time-dependent plasticity (STDP) mechanism, inspired by other models (Graupner & Brunel, 2012; Song et al., 2000). We first consider two bi-directionally connected neurons in a traditional STDP framework. Upon the occurrence of a spike event in a model neuron, post-synaptic weights are strengthened for any given pre-synaptic neuron that spiked beforehand or weakened in the vice versa condition.

The assumption being that the spike arriving at the post-synaptic connection must have either contributed to or competed with the spike event in question, depending on the directionality of the connection, leading to a reward or punishment of the synapse, respectively. To implement this, we here calculate potential synaptic plasticity via functions for long-term potentiation (F_LTP_) and long-term depression (F_LTD_) at the time of an eliciting spike (t) in Equations 4.1-4.2. Parameters were fitted to model the replication of data obtained from a hippocampal cell culture study (Huerta & Lisman, 1995) as shown in Figure 2, which also acts as a visual guide to the description of the set of Equations 4 & 5. Note that the measurement of global theta phase is dependent on the site of the observation, where it might be phase-reversed if measuring at the hippocampal fissure or at the cell body of CA3 (Hasselmo, 2005). As in prior models (Parish et al., 2018), we reverse theta phase from that of anatomical observations (Huerta & Lisman, 1995). This enables the functional selectivity and preferential binding of neurons that fire together out of phase, at a time when most cells are inhibited and therefore inactive.

**Figure 2.**
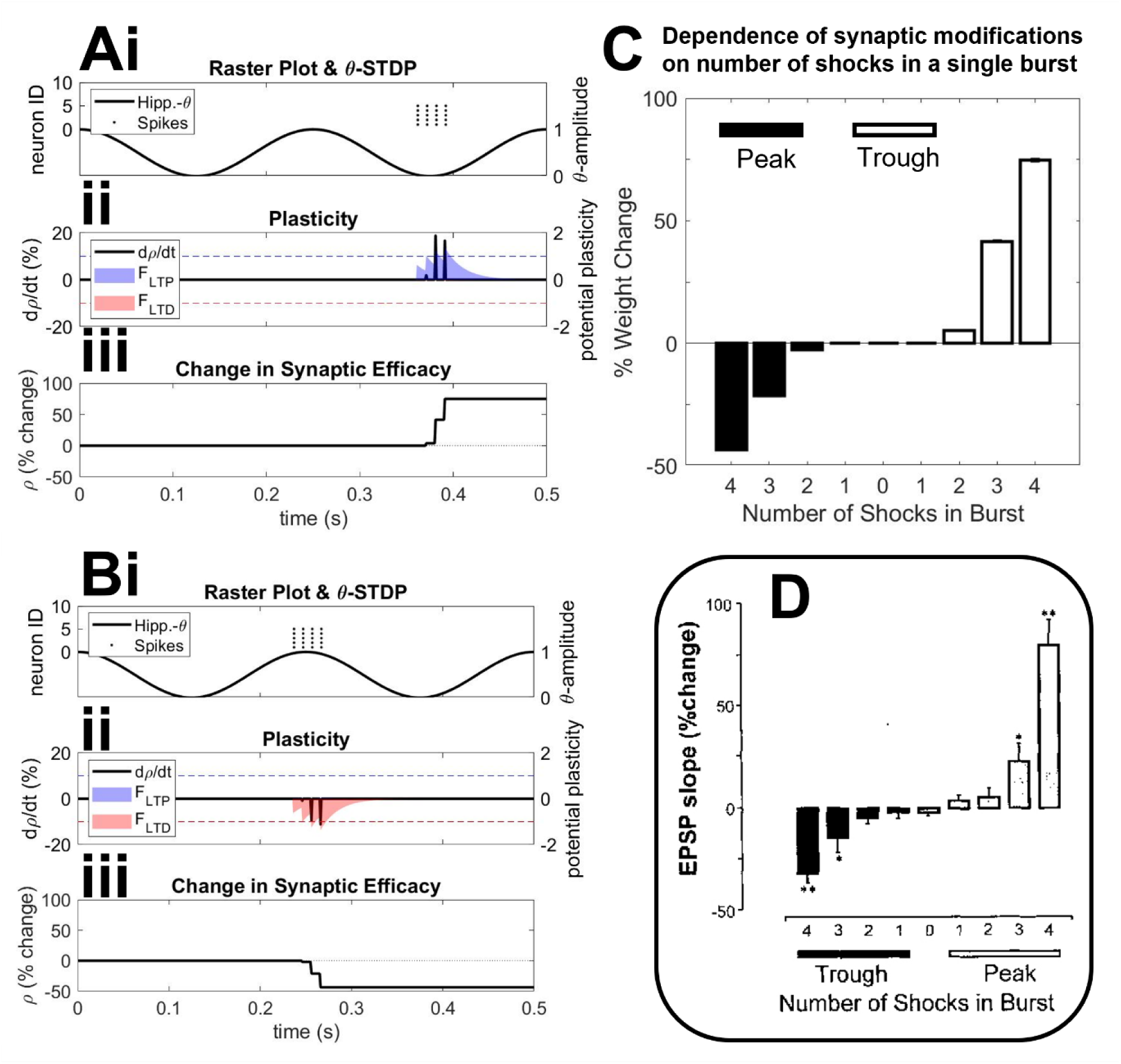
Evaluation of theta modulated spike-time dependent plasticity (STDP). ***A&B,*** independent simulations depicting synaptic plasticity after a single burst of 4 spikes at 100Hz (amplitude = 3pA), both in the trough ***(Ai)*** or the peak ***(Bi)*** of on-going theta oscillations. Potential plasticity induced by spike-pairings is calculated via functions (***A***&***Bii*** shaded regions; see Equations 4.1-4.2) for long-term potentiation (LTP; blue) & long-term depression (LTD; red). At the time of spike event, synapses undergo potentiation or depression (black lines; see Equations 5.1-5.2) if potential plasticity is above or below a potentiation or depression threshold (blue & red dotted lines, respectively). Overall synaptic change is calculated as a percentage to a baseline period through time ***(A&Biii)***. ***C,*** dependence on bursting for inducing plasticity, where the number of spikes in a burst was increased from 1-4 either in the peak or trough of ongoing theta (simulating 25 trials per condition). Overall synaptic change was calculated as the percentage difference to a baseline period. ***D***, earlier experimental observations that indicated the importance of bursting for inducing plasticity (Huerta & Lisman, 1995). Notation of “trough” & “peak” is dependent on the location of the recording site that describes theta phase. We prefer to flip this notation in relation to prior studies, to make clear the functional role theta might play in neuronal selectivity.

In the case of potentiation (Equation 4.1), potential LTP at the post-synaptic connection (i) is calculated as the summation of historic pre-synaptic spikes (n_pre_) that occurred before the spike event in question (where t_i_ < t), weighted by an absolute value (A_+_ = 0.65). Contributions of pre-synaptic spikes were proportional to an exponential decay, thus favouring spikes that occurred close together in time (τ_s_ = 20ms). Potential potentiation of the synapse in response to eliciting spike events is shown in Figure 2Aii (right hand axis, blue shaded region). In the case of depression (Equation 4.2), potential LTD at the pre-synaptic connection (j) was similarly calculated as the summation of historic post-synaptic spikes (n_post_) that occurred before the spike event in question (where t_j_ < t), weighted by an absolute value (A_-_ = 0.65). Similarly, potential depression of the synapse in response to eliciting spike events is shown in Figure 2Bii (right hand axis, red shaded region).

Further to this classical STDP framework, we here modulate learning by on-going theta oscillations (0≤ θ ≤1), as in previous models (Hasselmo, 2005; Ketz et al., 2013; Norman et al., 2006; Parish et al., 2018) and as suggested by empirical evidence (Huerta & Lisman, 1995; Pavlides et al., 1988). As a continuation of previous modelling work (Parish et al., 2018), where both LTP & LTD were multiplied by the phase of theta, we then provided preferential phases of general plasticity to active neurons. Here, we develop the learning rule by splitting LTP and LTD into opposing phases of theta (as exemplified in Equations 4.1-4.2, where historic spike events were further weighted by either theta or the inverse of theta for F_LTP_ & F_LTD_, respectively, as indicated in Figure 2A-B by the relationship of potential potentiation and depression to the phase at which spike events occur relative to an ongoing theta cycle). This change was crucial for the model to replicate further experimental work (Clouter et al., 2017; Wang et al., 2018), where nuanced offsets in the phase of active stimuli were preferentially rewarded or punished due to this segregation of continuous time into phases of potentiation and depression. In this way, stimuli that synchronised in phase were rewarded and bound together, whilst those that dropped out of phase with one another were actively punished.

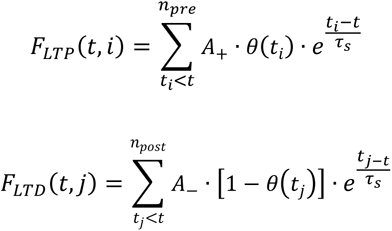

Equations 4.1 - 4.2 – Synaptic plasticity functions (F) calculate potential plasticity as the summation of the total number (n_post_ & n_pre_) of historic spike events (t_i_ & t_j_) arriving at a post-(i) or pre-synaptic (j) synapse relative to a given spike event (t), where an absolute value (A_+_/A_-_) is modulated by the difference in spike times and theta phase. Plasticity functions are separated for both long-term potentiation (LTP) & long-term depression (LTD), which are in turn modulated by opposing phases of a theta oscillation (0≤ θ ≤1).

For each spike event, Equations 5.1-5.2 were applied to all post- (i) & pre-synaptic (j) connections to the spiking neuron in a single use fashion, i.e. at the millisecond of spike occurrence (t) and not over a sustained period. The terms and parameters of Equations 5.1-5.2 were fit to model experimental observations of hippocampal cell cultures (Huerta & Lisman, 1995), and are visualised in the left-hand axis of Figure 2A-B (black lines). Synapses changed in proportion to their prior value (or the inverse of), such that 0 ≤ ρ ≤ 1. Synaptic change occurred at a constant rate for potentiation (γ_p_ = 1.5) and depression (γ_d_ = 0.75), where potentiation occurred at twice the rate of depression to more closely resemble theta modulated plasticity observed anatomically (Huerta & Lisman, 1995), as shown in Figure 2D. In these observations, the bursting of single cells was just as important in acting as a gateway to plasticity. This is captured in the usage of a Heaviside function (H[]), which provides either a 1 or a 0 dependent if potential plasticity (F_LTP_ or F_LTD_) was above a plasticity threshold (ε_LTP_ = 1 for potentiation & ε_LTD_ = 1 for depression), thus nullifying singlet or doublet spike pairings from triggering plasticity at the synapse. These thresholds are indicated by the dotted lines in Figure 2Aii & Bii (right-hand axis; blue for ε_LTP_, red for ε_LTD_). The amount of plasticity at the synapse was also moderated by the amount by which potential plasticity was above the plasticity threshold, causing a graded plasticity effect that exponentially increases the more spike pairings are contained within a burst (as indicated by experimental observations shown in Figure 2D). These alterations to the more traditional STDP rules that informed our model (Graupner & Brunel, 2012; Song et al., 2000), allowed us to closely replicate anatomical observations that synaptic plasticity is dependent on theta phase (Figure 2A-B) and bursting (Figure 2C) in hippocampal cell cultures (Huerta & Lisman, 1995). This in turn has allowed us to replicate more nuanced memory effects (Clouter et al., 2017; Wang et al., 2018) than the previous instantiation of the Sync/deSync model (Parish et al., 2018).

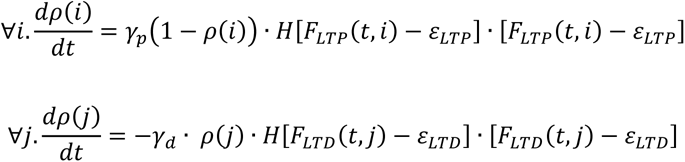

Equations 5.1 – 5.2 – Actual plasticity acting on a synapse (0≤ ρ≤1) at the occurrence of a spike event (t). Post- (i) & pre-synaptic (j) efficacy changes are induced by potential plasticity functions (F_LTP_ & F_LTD_, respectively) being above a threshold (ε_LTP_ & ε_LTD_, respectively). The sets i & j represent the indices of all post- & pre-synaptic synapses of the spiking neuron, respectively.

See also Supplementary Figure S1 for a more in-depth replication of the aforementioned anatomical observations (Huerta & Lisman, 1995). There we implement an additional rule to model the occurrence of observed hetero-synaptic plasticity on non-stimulated pathways (Equations S1-S2). With respect to Occam’s Razor principle, this was eventually excluded from the overall functionality of our model, as it did not directly influence results from our simulated paradigm and was also not central to the original anatomical observations. However, the additional rule might be useful to regulate network stability in the application of our theta-phase learning rule to large networks (Volgushev et al., 2016).

### NC system

Neurons within each subgroup of the neocortex (NC) had a 25% chance of being connected (W_max_ = 0.3). Connections of neurons between subgroups were not implemented in NC as it was assumed visual and auditory stimuli had not been previously associated. Synaptic plasticity (as described in Equations 4-5) was also not operating on cortical synapses as in the complimentary systems framework (O’Reilly et al., 2014) it is assumed that cortical plasticity occurs on a much slower timescale. Background noise for each NC neuron was estimated by Poisson distributed spike-trains (4000 spikes/s, W_max_ = 0.023). A cosine wave of 10 Hz (amplitude = 0.1pA) was fed into NC neurons via I_AC_. Two constant inputs were fed into each NC subgroup to simulate presentation of visual and auditory stimulus via I_DC_ (amplitude = 1.75pA). These inputs were either modulated by a cosine wave at different frequencies and phase offsets or not modulated depending on the simulation purpose, which will be provided below.

### Hippocampal system

The two subgroups of hippocampal neurons that represented visual and auditory stimuli were fully connected to their NC counterparts (Figure 1A; W_max_ = 0.35 for NC→Hip & W_max_ = 0.08 for Hip→NC synapses), as it was assumed both stimuli were previously known. Background noise for each hippocampal neuron was estimated by Poisson distributed spike-trains (1500 spikes/s, W_max_ = 0.015). A cosine wave of 4 Hz (amplitude = 0.25pA) was fed into each hippocampal neuron to model ongoing theta activity. Synapses within the entire hippocampus had a probability of 50% of forming a connection (W_max_ = 0.65), such that weights for intra-subgroup synapses were set to maximum and those for inter-subgroup synapses were initially set to 0. Synaptic plasticity was in effect on all hippocampal synapses, as described in Equations 4-5, allowing for the association of visual and auditory stimuli to take place within the hippocampus.

Critical to the model was the assumption of an intermediary relay node between NC and hippocampal subgroups, assumed to be located somewhere in the EC. We implemented a filter function to simplify the relay node as a filtered input, where the amplitude of the EPSP function (Equation 2) that was applied to spikes originating in the NC and arriving at hippocampal synapses was modulated via the pathway between EC and the hippocampus (Equation 6; W_EC_ = 0.3). This pathway is known to have a reversal phase relationship with hippocampal theta (Hasselmo, 2005), resulting in a stronger input at hippocampal theta trough and a weaker input at hippocampal theta peak. This intermediary filter allowed for model NC neurons that are entrained at a theta frequency to be active at and thus maximally induce plasticity in the appropriate hippocampal theta phase.

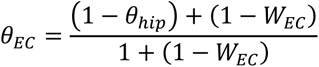

Equation 6 – Theta in the EC existed as the phase reversal of hippocampal theta (0≤ θ_hip_ ≤1), such that EC intermediary modulation of the EPSP function between NC and hippocampus neurons existed within the range 0→0.5≤ θ_EC_ ≤1, dependent on W_EC_.

### Simulation procedure and model evaluation

To compare our model with a human episodic memory paradigm (Clouter et al., 2017; Wang et al., 2018), we simultaneously fed two cosine waves (0≤ amplitude ≤1pA) into the visual and auditory NC subgroups (Figure 1B). A 2-second inter-stimulus interval was used before visual-auditory stimulus presentation. The two cosine waves were modulated at 4 Hz (theta), 1.652 Hz (delta), and 10.472 Hz (alpha) with auditory stimulus phase offsets of 0°, 90°, 180°, and 270° from the visual stimulus. The faster and slower frequencies were chosen to assess frequency specificity of the learning effect, as shown in Clouter et al. (2017). A baseline condition was also conducted by comparing learning in the theta 0° and 180° phase offset conditions with a non-oscillatory, constant input (no flicker condition in Clouter et al., 2017). To account for the difference in the amount of information between oscillatory and non-oscillatory conditions, the stimulus length in the no flicker condition was half of the length in the theta conditions (1.5 seconds).

To model the variability of brain activity entrained by an external rhythmic stimulus, we introduced noise that was formed by a normal distribution centred at each stimulus input frequency (i.e. 4 Hz, 1.652 Hz and 10.472 Hz) and with standard deviation (STD) of 0.015 multiplied by each input frequency, respectively, for additional independent simulations. Similarly, in another set of independent simulations (theta, delta, alpha and no flicker conditions), we introduced noise to hippocampal dynamics by randomising the hippocampal theta frequencies over a normal distribution centred at 4 Hz and with STD of 0.02, as well as randomising the EC phase offsets relative to ongoing theta with a normal distribution centred at 180° and with STD of 0.167. So far, for all simulations, ongoing alpha and theta cosine waves (i.e. I_AC_ in Equation 1) had a random phase at the beginning of each simulation. During learning, hippocampal theta phase was reset with a 180° offset from modulated visual stimulus input after stimulus onset (Figure 1B). Importantly, this enabled the fluctuation of theta modulated plasticity to be fully synchronised with stimulus inputs, thus maximising the learning potential of the model. Evidence for such a theta reset exists in several empirical studies (Rizzuto et al., 2003).

Model evaluation was done by comparing the full model with two compact versions of the model. In the theta-phase-learning only version, the STDP component was eliminated, and selected synapses were strengthened or weakened depending on hippocampal theta phase (i.e. removing the exponential components and redefining -1≤ theta ≤1 in Equations 4.1-4.2, as well as forcing the constants γp & -γd in Equations 5.1-5.2 to both be positive). This forced synapses to be bidirectionally and non-specifically weakened or strengthened at the inhibitory or excitatory phases of theta, respectively. In the STDP only version, we removed hippocampal theta activity, such that hippocampal theta phase did not reset with stimulus onset and both STDP weight changes and EPSP at synapses of input to the hippocampus were independent of ongoing theta phase, that is θ(t) was set to 1 in Equations 4.1-4.2 and θ_EC_ was set to 1 in Equation 6. This resulted in a lack of punishment for weak weight changes between groups with slightly overlapping time windows, a function previously performed by theta specific LTD. This caused sufficient learning in every phase offset condition to such an extent that they did not differ from each other (see Figure S2). To resolve this, we adjusted the range of input strength from between 0 and 1 to between -1 and 1 (amplitude = 1.75pA), thus more precisely defining the overlap of firing between two groups of neurons and punishing the weak weight changes between slightly overlapping groups, allowing us to evaluate the impact of STDP learning on the theta modulated inputs with different phase offsets. All simulations were run for 384 trials for each condition, which were then averaged across trials for each condition. For each simulation, we randomised a new set of initial synaptic connections as well as new Poisson distributed spike-trains for all conditions. We also initialised a new randomised frequency or EC phase offset for the simulations with noise.

## Results

### Simulated hippocampal weight change reproduces theta phase synchrony induced memory enhancement

We compared memory performance between the results of our model to those from human episodic memory studies that establish modulation of theta phase synchrony. Specifically, Clouter et al. (2017) and Wang et al. (2018) used a multisensory theta entrainment paradigm to externally manipulate the precise timing of visual and auditory inputs. As depicted in Figure 1B, luminance and amplitude of visual and auditory stimuli were modulated at 4 Hz with different phase offsets between the two stimuli: 0° (in-phase), 90°, 180° and 270° (out-of-phase). Human participants’ recall accuracy was evaluated by presenting the auditory stimulus and requesting participants to select the associated visual stimulus with which it was previously paired during the encoding phase. The encoding phase was simulated by feeding two, 4 Hz cosine waves that had phase offsets at either in-phase (0°) or out-of-phase (90°, 180°, and 270°) for three seconds into two subgroups of NC neurons that represented visual and auditory cortical neurons. The hippocampal theta phase was reset to be 180° offset from the visual stimulus by stimulus onset during encoding so that the LTP phase was aligned with the visual stimulus input. The entire procedure was simulated 384 times for each phase offset condition and the results were then averaged across simulation runs per condition.

To evaluate the recall performance of the model, the hippocampal weights from the Auditory to the Visual subgroup after learning were averaged between 2.75 to 3 seconds (one theta cycle) after stimulus onset. As shown in Figure 3A, weights after stimulus onset increase significantly in the 0° phase offset condition as compared to all other (asynchronous) conditions. In the 0° phase offset condition increases in weights are rhythmic at 4 Hz. This is because both inputs are synchronised with the inhibitory phase of theta. Neurons of both groups rarely fire at the excitatory phase of theta (i.e. the LTD inducing phase), therefore weight changes are mainly positive until weights reach their maximum. This is also reflected by the hippocampal firing activity during learning for the group of neurons corresponding to visual inputs. After stimulus onset, an increase in firing of the visual neurons in the 0° phase offset condition is evident across time due to the weight change (Figure 3B), which leads to changes in the input current for I_syn_ (see Methods).

**Figure 3.**
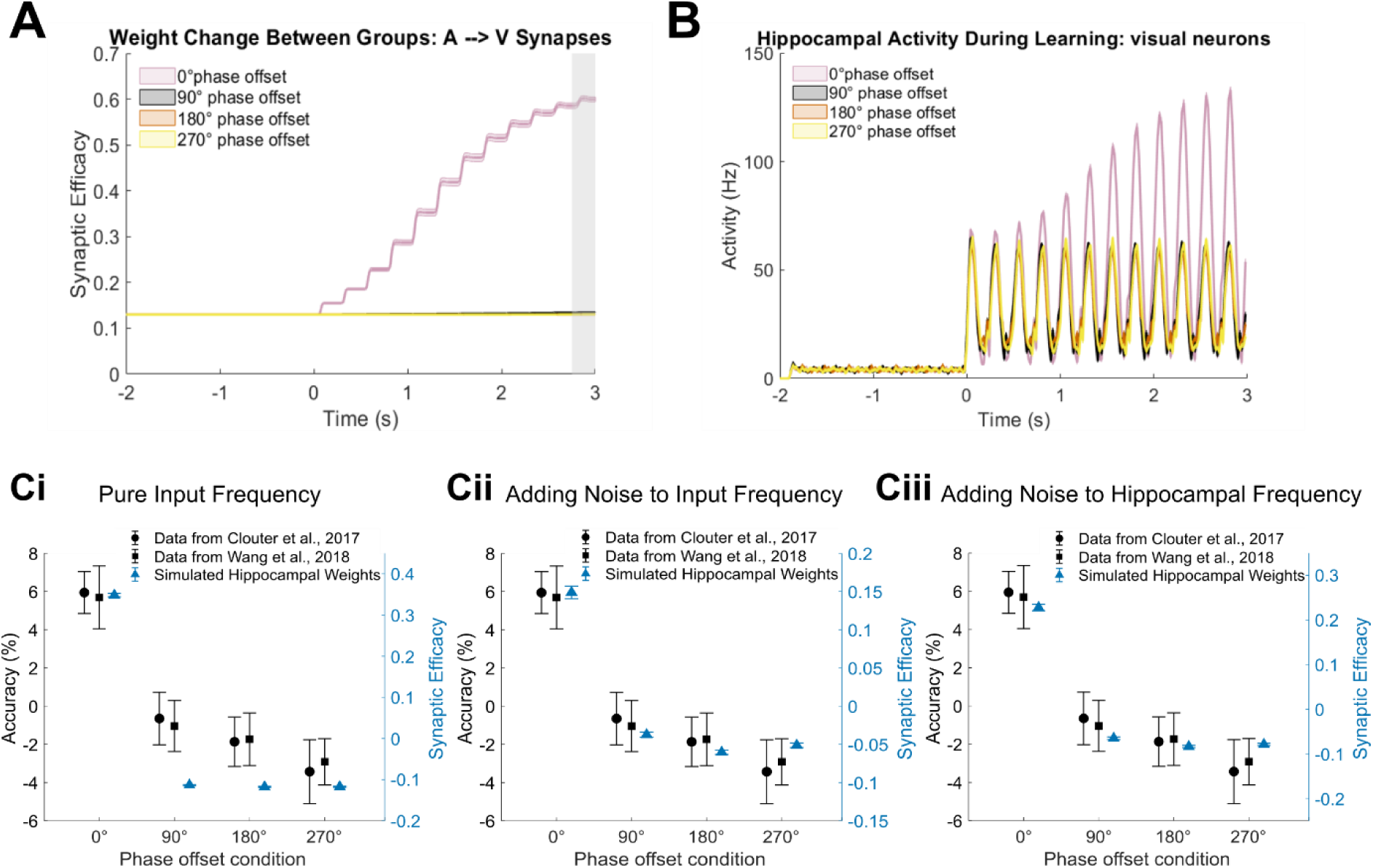
Recall performance of the model as a function of phase offset condition when stimulus inputs are modulated at theta frequency (4 Hz). *A,* Hippocampal weight change from the Auditory to the Visual subgroup during learning. Weights from the Auditory to the Visual subgroup increase significantly in the 0° phase offset condition after stimulus onset. The weights after learning are averaged between 2.75 to 3 seconds after stimulus onset (the grey shaded area). *B,* Firing rate of hippocampal visual neurons responding to the auditory stimulus during learning. *C,* Mean of weights from the Auditory to the Visual subgroup after learning from 384 simulations and empirical data from Clouter et al. (2017) and Wang et al. (2018). Accuracy is normalized by subtracting the mean over all 4 phase offset conditions to make the between studies data more comparable (i.e. to correct for differences in absolute memory performance between studies). *Ci,* simulations of pure input frequency, 4 Hz, hippocampal frequency, 4 Hz, and EC phase offset, 180° from hippocampal theta. *Cii,* simulations of input frequencies randomly drawn from normal distribution with mean 4, and standard deviation (STD) 0.015*4, pure hippocampal frequency, 4 Hz, and EC phase offset, 180° from hippocampal theta. *Ciii,* simulations of pure input frequency, 4 Hz, hippocampal frequencies randomly drawn from normal distribution with mean 4, and STD 0.02, and EC phase offsets randomly draw from normal distribution with mean 180°, and STD 0.167. Error bars represent standard error (SE).

Figure 3Ci shows mean weights simulated for each phase offset condition compared with experimental data from Clouter et al. (2017) and Wang et al. (2018). Consistent with both experimental data sets the model shows highest memory performance for the 0° phase offset condition relative to the other three out-of-phase conditions. Moreover, the model successfully replicates the previously observed pattern that memory performance in the out-of-phase conditions (90°, 180°, and 270°) did not differ from each other. The fact that the recall performance in the 90° and 270° phase offset conditions does not differ from the performance in the 180° phase offset condition could be caused by the interaction between theta-phase-dependent and STDP learning. Inputs of auditory neurons on visual neurons in the 90° and 270° phase offset conditions are less synchronised with the LTP phase within a theta cycle, thus the synaptic changes are weighted less by corresponding theta phases. As synaptic weight changes decay exponentially across time, the resultant small weight changes will decay drastically when the next visual stimulation peak induces substantial spiking events, given that the delay between auditory and visual stimulation peak is longer than 50 ms.

We note the difference, however, between the modelled and empirical data, whereby the model shows relatively larger differences between the synchronous and asynchronous conditions compared with the experimental data. This could be because this version of the model had zero noise, i.e. it assumed a perfect transmission of rhythmic activity from the sensory channels to the brain, as well as a perfect alignment of the hippocampal theta rhythm to the sensory input. Both are unlikely to be the case in the human brain, which is expected to show trial-by-trial variability to rhythmic sensory input (see Wang et al., 2018). To model variability of sensory transmitted rhythms, we generated a normal distribution that centred at 4 Hz, with STD of 0.015 multiplied by 4 as noisy input frequencies (see Methods).

Incorporating such noise sources led to a better fit of the simulated data to the empirical data (Figure 3 Cii). Additionally, we modelled variability of hippocampal theta by varying theta frequency (i.e. using a normal distribution centred at 4 Hz with STD of 0.02). In addition, the EC phase offsets relative to ongoing theta was also randomised with a normal distribution centred at 180° with STD of 0.167. Again, the simulation results showed a more similar pattern as observed in the empirical studies (Figure 3 Ciii).

### Simulated hippocampal weight change reproduces theta specificity of the phase synchrony induced memory enhancement

We tested whether our model would also show theta specificity of the phase offset effects, as revealed by Clouter et al. (2017). More specifically, Clouter et al. showed that the difference in memory performance between synchronous (0°) and asynchronous (90°, 180°, 270°) conditions is specific to theta stimulation and is not observed for a slower (1.7 Hz) or faster (10.5 Hz) frequency. We therefore fed two cosine waves, modulated at same phase offsets, and modulated their frequencies at 1.7 Hz (delta) and 10.5 Hz (alpha). This was done to prevent the chosen frequency bands occurring in a harmonic relationship with 4 Hz (Pletzer et al., 2010). To compare these results with the data from Clouter et al., we averaged simulated hippocampal weights after encoding across the three out-of-phase conditions (90°, 180° and 270°) to yield a single asynchronous measure (Figure 4Aii). The modelled results replicated the pattern observed by Clouter et al. (2017) in showing that the memory difference between synchronous and asynchronous conditions is significantly greater at theta compared to the other two frequencies (Figure 4Ai and 4Aii). Moreover, the model replicates the findings of Clouter et al. in showing that memory performance in the synchronous condition at theta is better than in the same conditions of the two control frequencies. This suggests that synchronous stimulation improves memory specifically in theta frequency, reflecting coordinated timing of external inputs relative to hippocampal theta. Given that the alpha- and delta-modulated inputs do not coincide with the LTP phase during learning, firing could lead to LTP or LTD or, alternatively, half-weighted LTP and LTD (e.g. Figure 1A). Therefore, only synchronisation at the hippocampus’ preferred frequency is more likely to induce effective associative learning.

**Figure 4.**
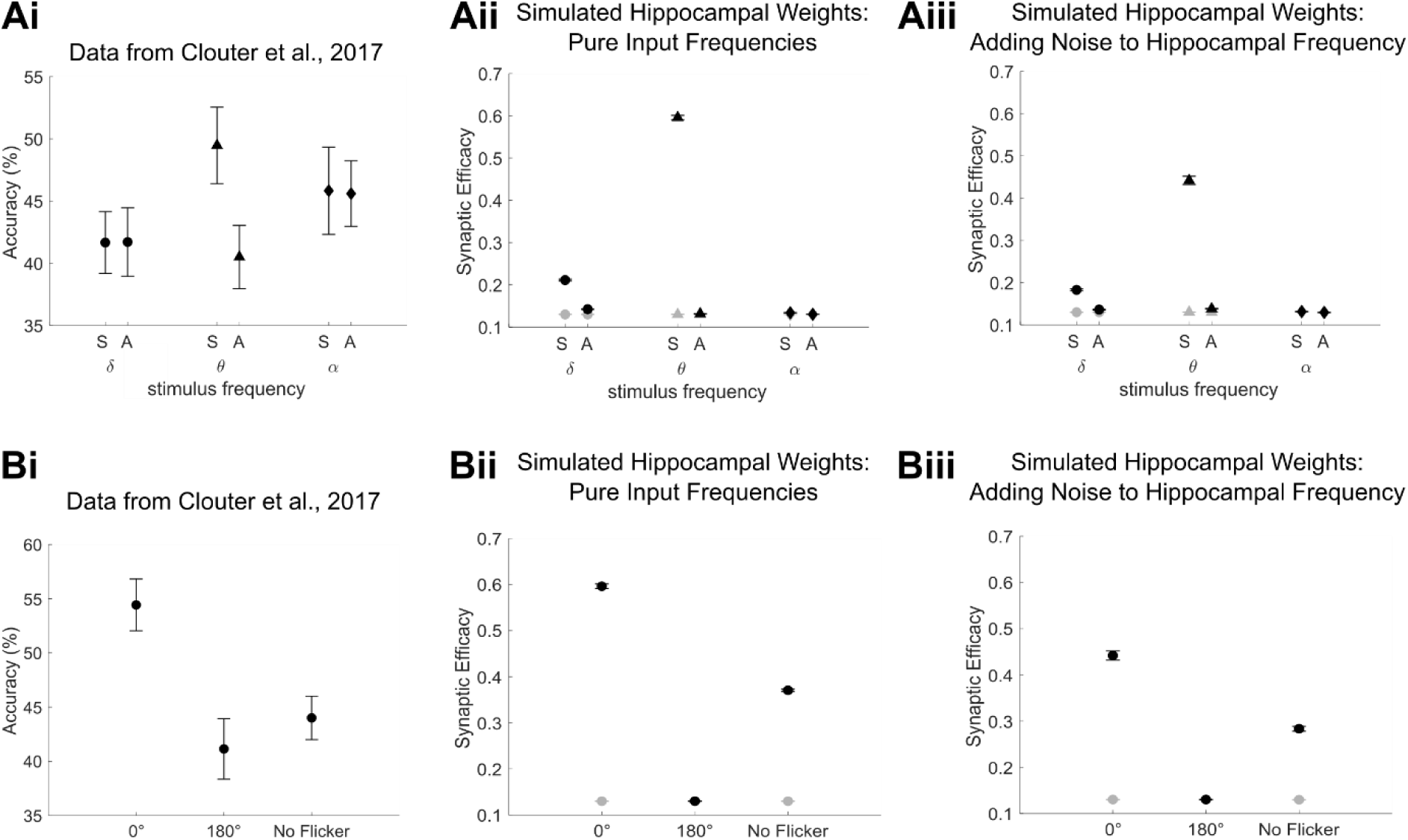
Recall performance as a function of degree of phase synchronisation in theta modulated condition and other control conditions. ***Ai,*** Data from Clouter et al. (2017) showing recall accuracy when the movies and sounds were flickering in synchrony (S) or out of synchrony (A) at delta, theta and alpha frequencies. ***Aii & Aiii,*** Mean of weights from the Auditory to the Visual subgroup after learning from 384 simulations. ***Aii,*** simulations of pure input frequencies, delta: 1.7 Hz, theta: 4 Hz, alpha: 10.5 Hz, hippocampal frequency, 4 Hz, and EC phase offset, 180° from hippocampal theta. ***Aiii,,*** simulations of pure input frequency, 4 Hz, hippocampal frequencies randomly drawn from normal distribution with mean 4, and STD 0.02, and EC phase offsets randomly drawn from normal distribution with mean 180°, and STD 0.167. ***Bi,*** Data from Clouter et al. (2017) showing recall accuracy when the movies and sounds were presented at 0-degree phase offset, 180-degrees phase offset, or were unmodulated. ***Bii & Biii,*** Mean of weights from the Auditory to the Visual subgroup after learning from 384 simulations. ***Bii,*** simulations of pure input frequencies, hippocampal frequency, 4 Hz, and EC phase offset, 180° from hippocampal theta. ***Bii,*** simulations of pure input frequency, 4 Hz, hippocampal frequencies randomly drawn from normal distribution with mean 4, and STD 0.02, and EC phase offsets randomly drawn from normal distribution with mean 180°, and STD 0.167. All error bars represent SE. Black dots represent hippocampal weights after learning. Gray dots represent hippocampal weights averaged between -1.75 s and 0 during pre-stimulus baseline.

Our model also replicates findings from another control experiment of Clouter et al. showing that flickering visual and auditory stimuli at 4 Hz in synchrony boosts memory beyond a natural non-flickering condition (Figure 4Bi). To simulate this, two constant stimulus inputs were fed into NC auditory and visual neurons during encoding. The length of the inputs was reduced to 1.5 seconds to control for the amount of overall stimulus presentation time in the flickering condition (i.e. the screen was essentially blank for half of the time due to the flicker). Figure 4Bii reveals that simulated hippocampal weights after learning are higher in the 0° phase offset condition than in either the 180° phase offset condition or the non-flickering condition. Because of the role of EC modulation in filtering NC inputs to be more active in the theta LTP phase and less active in the theta LTD phase (Methods), the constant inputs result in substantial associate learning as compared to the baseline. However, they are still less optimal than the theta synchronised inputs, which is consistent with the empirical data, confirming that the theta synchrony enhanced memory effect is indeed a result of maximally optimising input timing relative to the hippocampal theta oscillation. Figures 4Aiii and 4Biii show that the patterns still hold after adding noise into hippocampal dynamics. Moreover, the patterns resemble the empirical data better than before adding noise, which is reflected in a smaller difference between the theta synchronous condition and the other conditions.

### Theta-phase-dependent and STDP learning rules are necessary to reproduce the theta-induced memory effect

We compared the recall performance of the model to two alternative versions. The first is a model of purely theta-phase-dependent learning. For this version we eliminated the STDP component such that synaptic weights are modified solely depending on hippocampal theta phase. For example, if a hippocampal neuron from the auditory subgroup fires during the inhibitory (i.e. LTP inducing) phase, synapses between this neuron and its connected visual neurons will be potentiated, whereas the synapses between the neuron and its connected visual neurons will be depressed if it fires during the excitatory phase (i.e. LTD including phase). Hippocampal weights for the four phase offset conditions were fit to the two empirical datasets (Clouter et al., 2017; Wang et al., 2018) using linear least squares. Figures 5Ai and 5Bi show that the theta-phase-dependent learning only model does not fit the empirical data as well as the full model. The aforementioned result is likely due to the fact that memory performance for the theta phase only model increases depending on the degree to which the inputs overlapped with theta LTP phase. Weights in the 90° and 270° phase offset conditions are therefore slightly better than in the 180° phase offset condition but worse than in the 0° phase offset condition. We conducted F-tests to statistically compare the fitting of the theta phase only model with the fitting of the full model. The residual sum of squares (RSS) for the theta only version was significantly larger than the RSS for the full model (Fitting to Clouter et al. (2017) theta-phase-learning only vs. full: *F* (1, 3) = 17.11, *p* < 0.05; Fitting to Wang et al. (2018) theta-phase-learning only vs. full: F (1, 3) = 33.90, p < 0.05). We suggest this outcome arises because learning is cancelled in the 180° phase offset condition, as the auditory inputs cause firing at theta LTD phase, whereas firing at 90° and 270° lead only to a reduction in the positive weights, as compared to firing exactly at theta LTP phase (i.e. 0°).

**Figure 5.**
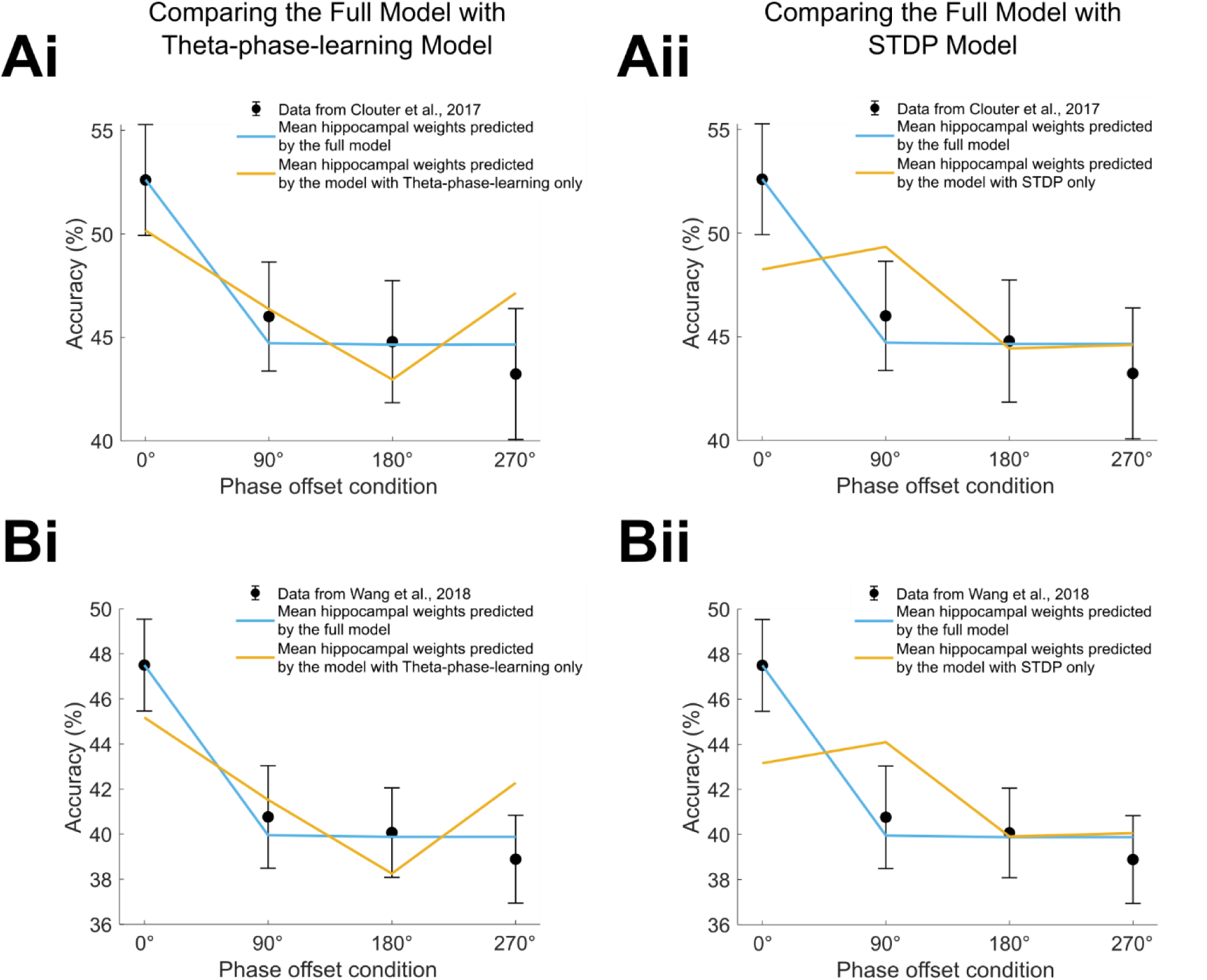
Model comparisons in recall performance between the full model and two alternative versions. ***A,*** Mean of weights from the Auditory to the Visual subgroup after learning simulated by two versions of model was fit to the data from Clouter et al. (2017). ***Ai,*** The full model was compared with a theta-phase-learning only version of the model. ***Aii,*** Same as Ai, but the comparison was between the full model and an STDP only version of the model. ***B,*** Same as ***A***, but mean of weights from the Auditory to the Visual subgroup after learning simulated by two versions of model was fit to the data from Wang et al. (2018). All simulations were done with pure input frequency, 4 Hz, hippocampal frequency, 4 Hz, and EC phase offset, 180° from hippocampal theta. All error bars represent SE.

Next, we created a version of the model i.e., the STDP only model, where all elements relating to theta modulated plasticity were removed. In this version of the model, weight changes can occur at any time, completely independent of theta phase, if spike timing between a sending neuron and receiving neuron is within a short time window (< 50 ms). This means that if enough spikes overlap in time between inputs, learning strength will not differ between phase offset conditions. To enlarge the spike timing gap between two inputs in each phase offset condition so the impact of STDP alone can be evaluated, we adjusted the range of the 4 Hz cosine wave to be between -1 and 1 which effectively narrows the time window for spiking. This leads to the largest gap in firing between two groups of neurons in the 180° phase offset condition because there is no overlap in spiking between the two groups. A few spikes overlapped in the 90° and 270° phase offset conditions, but the difference is that the spikes in the auditory group of the 90° condition are leading the spikes in the visual group, whereas in the 270° condition, the neurons of auditory group firing is lagging the visual group. Therefore, LTP in the synapses from auditory to visual groups is expected to occur in the 90° phase offset condition. In contrast, LTD should mainly occur in the synapses from auditory to visual groups in the 180° and 270° conditions. This is confirmed by Figures 5Aii and 5Bii that show enhanced memory performance in the 90° phase offset condition, whereas recall performance decreases exponentially from the 90° phase offset condition to the 180° and 270° phase offset conditions, with memory performance not being different between 180° and 270° phase offset conditions. However, memory performance is also enhanced for the 0° phase offset condition, where memory performance appears slightly lower compared to the 90° condition. This might be due to the complete overlap in neurons firing between two groups in the 0° condition. Since theta phase does not modulate weight changes anymore, such complete overlapping would lead to a reward and punishment simultaneously, hence balancing at a similar level in weights as in the 90° condition. As a result, the model with STDP learning mechanisms produces only a poor fit to the empirical data compared to the full model. F-tests showed that RSS for the STDP only version is significantly larger than the RSS for the full model (Fitting to Clouter et al. STDP only (input range -1 and 1) vs. full: *F* (1, 3) = 23.08, *p* < 0.05; Fitting to Wang et al. STDP only (input range -1 and 1) vs. full: *F* (1, 3) = 52.69, *p* < 0.05). Therefore, implementing both STDP and theta-phase-dependent learning mechanisms is essential to replicate the theta-induced memory effect from human episodic memory experiments.

## Discussion

We present a simple model involving a hippocampal system that implements STDP learning, where LTP and LTD are modulated by opposing phases of an ongoing theta oscillation. The model reproduces important findings from a hippocampal cell culture study showing that a brief burst of a few spikes at opposing theta phases induces LTP and LTD, respectively (Huerta & Lisman, 1995). Using parameters fitted to reproduce these rodent data, we can replicate several key findings from human episodic memory studies (Clouter et al., 2017; Wang et al., 2018). The simulated hippocampus received inputs from two different groups of neocortical neurons, visual and auditory neurons that are modulated at the same frequency as the hippocampal theta frequency. Synchronising the inputs to be in-phase with the fluctuation of theta-phase-dependent LTP induces more effective hippocampal synaptic connectivity, as compared to when the inputs are desynchronised. Such learning effects induced by phase synchronisation were only present when the inputs were modulated at the same frequency as hippocampal theta but not at alpha or delta frequencies that do not modulate hippocampal LTP or LTD. Moreover, our model replicates the result that synchronizing input at theta frequency improved memory compared to unmodulated input, suggesting that the theta phase induced memory effect is highly attributed to the hippocampal dynamics rather than purely perceptual binding (Clouter et al., 2017). To simulate the exact pattern of recall accuracy observed in the human episodic memory studies, the hippocampus must have both synaptic modification mechanisms. On one hand, the theta-phase-dependent learning only model fails to replicate the pattern that learning in the 90°, 270° and 180° conditions did not differ from each other, as these conditions seemingly benefit from the additional subtlety that STDP brings with regards to slight differences in spike timing. On the other hand, STDP learning alone shows that learning only depends on pre-synaptic and post-synaptic inputs timing. Therefore, both the 0° and 90° phase offset conditions show enhanced learning, as compared to the 180° and 270° conditions, hence failing to reproduce the empirical findings.

Theta-phase-dependent plasticity as implemented in our model separately modulates the two components of STDP, LTP and LTD. Such modulation by opposing theta phases is inspired by several theta learning models. Hasselmo et al. (2002) and Ketz et al. (2013) modelled theta phase reversal between hippocampal subregion pathways as key functional processes in memory formation. In Hasselmo et al.’s model, phase differences between EC and hippocampal CA1 either encourages LTP within hippocampus CA3 to CA1 in an effective encoding phase or blocks this activation pathway in favour of encouraging cortical activation through the reverse pathways in an effective retrieval phase. Thus, theta phase reversal efficiently separates encoding and retrieval, enabling pattern separation of overlapping memories. Ketz et al. (2013) modelled similar hippocampal theta dynamics, additionally showing that theta phase learning is more effective for error-driven learning, as compared to purely Hebbian learning, thus showing how theta can further hone accurate memory formation in an iterative manner. Another model in this line of thought operates at a more theoretical level with the aim of solving some issues arising from competing and overlapping representations in neural network learning (Norman et al., 2006). Rather than modulate specific neuronal pathways, the model implements general oscillatory inhibition. Norman et al.’s model is able to solve competition between overlapping memories by showing how an increase in inhibition can strengthen target memories, whilst a decrease in inhibition can weaken competitors. Thus, theta, which Norman et al. (2006) hypothesise to be the candidate mechanism for modulating network stability in this way, can efficiently perform pattern separation on a population simply via distinct phases for excitation and inhibition. Whilst our model does not focus on the role of theta phase in recurrent medial-temporal-lobe pathways, nor on some key hippocampal functionality such as pattern separation and error-driven learning, we are able to show how preferential stimulus encoding can be achieved through the extrapolation of theta induced effects at the neuronal level alone, i.e. when overlapping sensory input is in-phase with theta modulated LTP then weights are stronger than when input is out-of-phase with these LTP fluctuations. We do this by marrying the notion of reward and punishment from STDP dynamics with the theta dynamics from the Hasselmo et al. (2002) and Norman et al. (2006) models, where distinct phases of theta provide contrasting functional windows that are vital for continued network stability when creating new memories. By exploring the emergent network properties that arise from these changes at the cellular level, we hope to show the multi-faceted role that the theta frequency might play in memory formation, from both a network and cellular perspective.

This model is a continuation of the previous Sync/deSync model (Parish et al., 2018). In the Sync/deSync model, though LTP and LTD is modulated by opposing phases of the ongoing theta rhythm, LTD happening at the theta peak is applied by a global passive decay. The passive decay is an exponential function that is multiplied by the complement of theta phases, which leads to maximal weight decay at theta peak and 0 at theta trough. The current model replaces this with an active theta-phase-specific LTD, which is crucial to replicate the key findings from Clouter et al. (2017), namely that the phase synchrony induced memory advantage is specific to theta modulated stimulus inputs. When stimuli are modulated at non-hippocampally preferred frequencies (i.e. outside of theta), neuronal firing at the theta LTD phase is more likely to happen, and less likely to happen at the LTP phase. The active theta-phase-specific LTP and LTD thus allows more potentiation of synapses when stimulus inputs are theta modulated and in synchrony, and more LTD when stimulus inputs are modulated with delta and alpha frequencies, even though they are in synchrony. This in turn allows the Sync/deSync model to be much more sensitive to very slight differences in the timings of concurrent stimuli, which was not an immediate concern during its original conception as a first proof of principle model that related oscillations to human episodic memory formation.

In rodents, LTP and LTD can be induced by burst stimulation at opposing hippocampal theta phases, respectively (Hölscher et al., 1997; Huerta & Lisman, 1995; Hyman et al., 2003; Pavlides et al., 1988). In humans, theta phase synchronisation can enhance episodic memory performance (Clouter et al., 2017; Wang et al., 2018). Our model can account for data from both rodents and humans, which connects human episodic memory behaviour to synaptic modifications induced by non-neurophysiological stimulation in the animal brain. Our model provides an explanation to the findings in human electrophysiological studies, which show that hippocampal theta synchronization supports episodic encoding, especially when there is need to form arbitrary associations (Backus et al., 2016; Kota et al., 2020; Staudigl & Hanslmayr, 2013). Though those findings are correlational rather than causal, our model suggests that theta synchronization links to stronger synaptic weights between corresponding hippocampal neurons after encoding. Our model predicts that to induce substantial LTP in synapses between hippocampal neurons, cortical inputs must be in-phase with the fluctuation of LTP. In contrast, if any input is desynchronised with the oscillatory changes of LTP, synaptic weights are very weak between the hippocampal neurons corresponding to those inputs. The phase of hippocampal theta oscillation is reset to coordinate the timing of inputs with time windows of LTP, which is a key characteristic in our model to optimise learning. Indeed, hippocampal theta phase resetting has been found to occur after stimulus onset during memory encoding (Rutishauser et al., 2010). Further, Kota et al. (2020) shows in an iEEG study that ongoing hippocampal theta oscillation phase was reset to support episodic encoding. Such phase reset indicates a mechanism for inputs to be more likely to undergo LTP (McCartney et al., 2004; Mormann et al., 2005; Rizzuto et al., 2003). Moreover, a rodent study recently reveals that using visual rhythmic stimulation at gamma frequency (40 Hz) can entrain gamma activity in the hippocampus and preserve hippocampal neurons and synapses (Adaikkan et al., 2019). Our model suggests this could also happen in the human hippocampus, where theta phase is reset by sensory stimuli, thus aligning stimulus inputs to be more likely to induce LTP and form associations.

Investigating the cellular mechanisms of human learning and memory is challenging as it is relatively difficult to access human single neurons *in vivo*. Although there is evidence on STDP in the human hippocampus *in vitro* showing that LTP was induced in a wider time window as compared to rodent hippocampal STDP (Silva et al., 2010). It is still unclear whether the same STDP rule applies in the human hippocampus *in vivo*. In our model, the classical STDP rule observed in the rodent hippocampus is implemented (Bi & Poo, 2001; Parish et al., 2018; Song et al., 2000). Our model efficiently forms new associations in a one-shot manner as required by episodic memory. In human single neuron studies, neurons firing selectivity in the Medial Temporal Lobe (MTL) was extended to the learned, associated contextual pictures or temporal sequences (Ison et al., 2015; Reddy et al., 2015). Synaptic plasticity is very likely to be the circuit mechanism supporting episodic formation, which provides evidence that neuronal mechanisms of memory formation might be universal across species. However, our model predicts that STDP alone will result in similar performance between 0° phase offset and 90° phase offset conditions, since LTP and LTD can happen at the same time if spikes are completely overlapped. Therefore, the two learning mechanisms, theta-phase-dependent plasticity and STDP are both essential to reproduce the empirical findings. They might interact to support episodic binding. STDP is a passive learning process so that weight changes happen at any time. However, if combined with theta-phase-dependent learning, an active learning mechanism, the model accurately reproduces the empirical data. This is consistent with the notion that theta oscillations might compress the neural events that have happened on longer time scales, thus enabling STDP in action (Hanslmayr et al., 2016). In the rodent hippocampus, the LTP component of STDP is dependent on spike timing triggered during a theta oscillation, but not during a low frequency activity (Wittenberg & Wang, 2006). This is consistent with our model, which shows that the learning effect is specific to theta-modulated inputs and all other conditions modulated by non-theta frequencies showed similarly low performance. Interestingly, a recent intracranial EEG study showed that in the human hippocampus, co-firing of neurons at 20 – 40 ms indicated successful episodic memory formation, while co-firing at longer delay of 60 ms resulted in forgotten triplets. Moreover, such subsequent memory related co-firing effect was specific to the neuron pairs that were coupled to distal theta and local gamma oscillations. Reversal of the order of the neuron pairs resulted in an LTD-like effect, where a shorter delay resulted in triplets subsequently forgotten (Roux et al., 2021). In the human hippocampus, the interaction of the two learning mechanisms, STDP and theta-phase-dependent plasticity, likely provides an explanation for the hippocampus’ role in actively coordinating the timing of input arrival, hence binding the inputs into long-term memory.

We simulated our data by matching the stimulus input frequency with the hippocampal theta frequency. Our empirical data showed that there was variability in sensory entrainment. We therefore modelled variability of input frequencies as well as hippocampal theta dynamics. Unsurprisingly, such variability weakened synaptic weights. This variability might come from attentional modulation (Müller et al., 2006; Tiitinen et al., 1993) or neocortex-hippocampus feedback loops where hippocampal theta rhythms entrain the neocortex to allow more optimal information transmission from the neocortex to the hippocampus (Sirota et al., 2008). This could be tested by directionality analyses in further experimental work using the same paradigm and recording hippocampal and cortical activity, or by implementing additional modules in future models. Moreover, future work might benefit from detecting individual hippocampal theta frequency, then matching the stimulus input frequency with the hippocampal theta to optimise the theta synchrony induced memory effect.

In conclusion, our model successfully reproduced results in the human episodic memory studies that show a causal role of theta phase synchronisation in episodic memory formation. Our findings provide new and important computational evidence for a combined role of two well-known synaptic plasticity mechanisms to modulate theta phase dependent memory effects in humans (i.e. STDP and theta phase dependent plasticity).

## Acknowledgements

This research was supported by grants from the European Research Council (https://erc.europa.eu/, Consolidator Grant 647954; to S.H.); Wolfson Foundation and Royal Society (https://royalsociety. org/grants-schemes-awards/grants/wolfson-researchmerit/; to S.H.); The Economic Social Sciences Research Council (https://esrc.ukri.org/, ES/R010072/1; to S.H. and K.S.).

## Competing interests

The authors declare that no competing interests exist.

## Data and Code availability

All human data and the Matlab code of the model to run the simulations can be downloaded at: http://github.com/GP2789/Sync-deSync-model-TIME.

---

**Figure S1.**
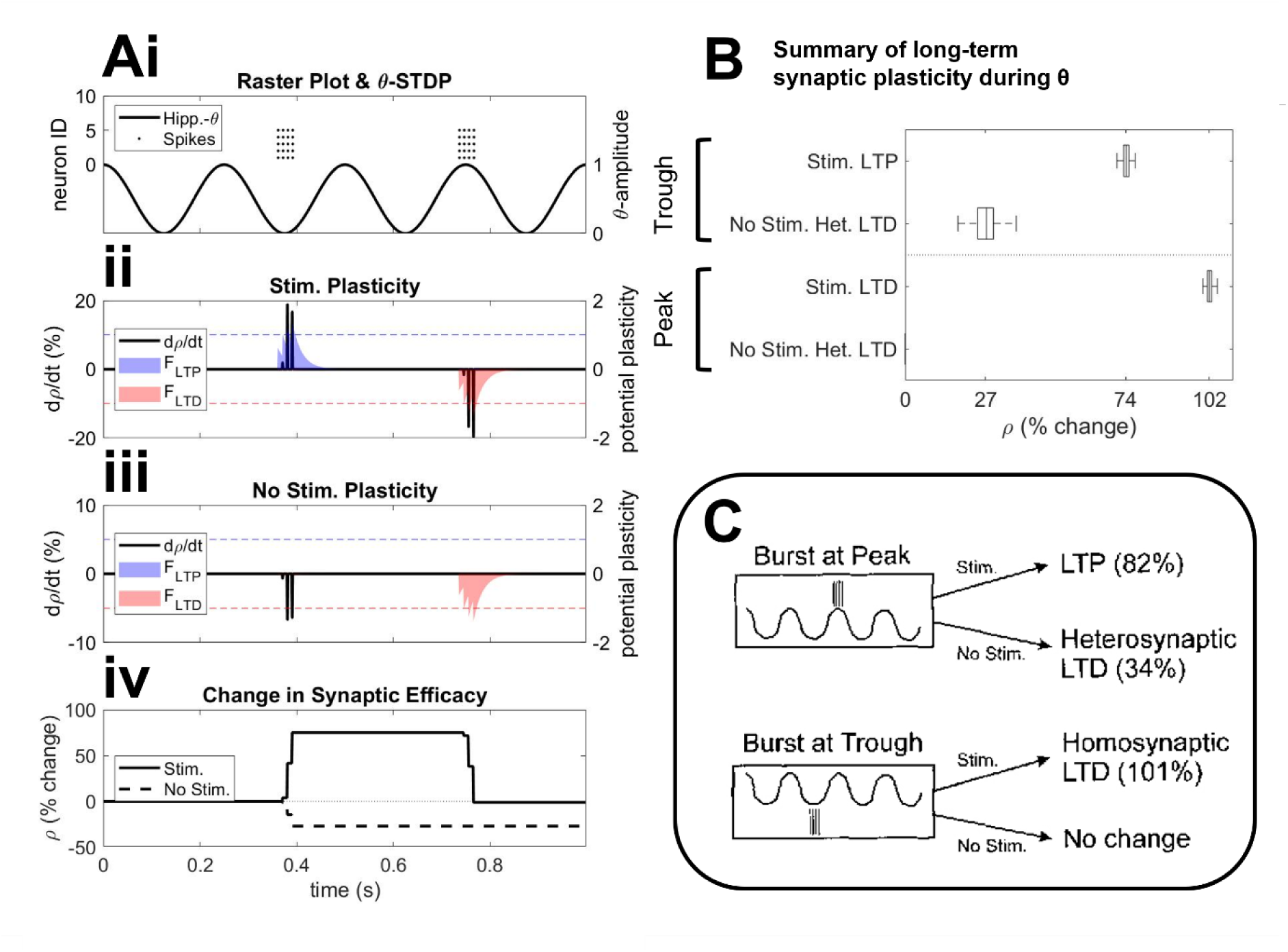
Summary of long-term synaptic plasticity during theta. ***Ai-iv,*** a single simulation depicting synaptic plasticity after two bursts of 4 spikes at 100Hz (amplitude = 3pA), first at the trough and then at the peak of on-going theta oscillations ***(Ai)***. Potential plasticity induced by spike-pairings is calculated via functions (***Aii-iii*** shaded regions; see Equations 4.1-4.2) for long-term potentiation (LTP; blue) & long-term depression (LTD; red). At the time of spike event, synapses undergo potentiation or depression (black lines; see Equations 5.1-5.2) if potential plasticity is above or below a potentiation or depression threshold (blue & red dotted lines, respectively). Overall synaptic change is calculated as a percentage to a baseline period through time ***(Aiv)***. For this simulation only, we here make use of an additional equation that induces heterosynaptic LTD on non-stimulated (No Stim.) pathways ***(Aiii)***, that was proportional to the amount of LTP that occurred on stimulated pathways (Stim.) (see Equations S1-S2). ***B***, a summary of synaptic plasticity during theta, where confidence bars indicate variability over 100 trials. The percentage for LTP on stimulated pathways (Stim. LTP) and heterosynaptic LTD on unstimulated pathways (No Stim. Het. LTD) is the absolute of the percentage change of synapses relative to a baseline period. The percentage for LTD on stimulated pathways (Stim. LTD) is the depression produced by a single burst compared to the potentiation that preceded it (e.g., 100% is complete reversal). ***C***, experimental observations (Huerta & Lisman, 1995) surmising the relationship of theta and neuronal activity.

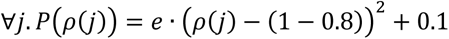

Equation S1 – Probability of synaptic plasticity on any pre-synaptic connection (j) of a spiking neuron where post-synaptic long-term potentiation (LTP) has taken place.

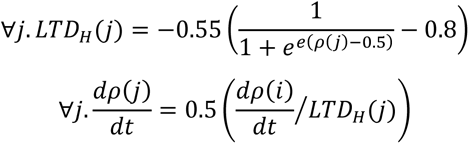

Equations S2.2 – 2.3 – Heterosynaptic long-term depression (LTD_H_) acting on any pre-synaptic connection (j) that has met the probability for change in Equation S1. The amount of heterosynaptic LTD is further normalised against the amount of prior LTP on the post-synaptic connection (dρ(i)/dt from Equation 6.1), such that the sum of LTD on pre-synaptic connections is roughly equal to half the amount of LTP that occurred on post-synaptic connections.

**Figure S2.**
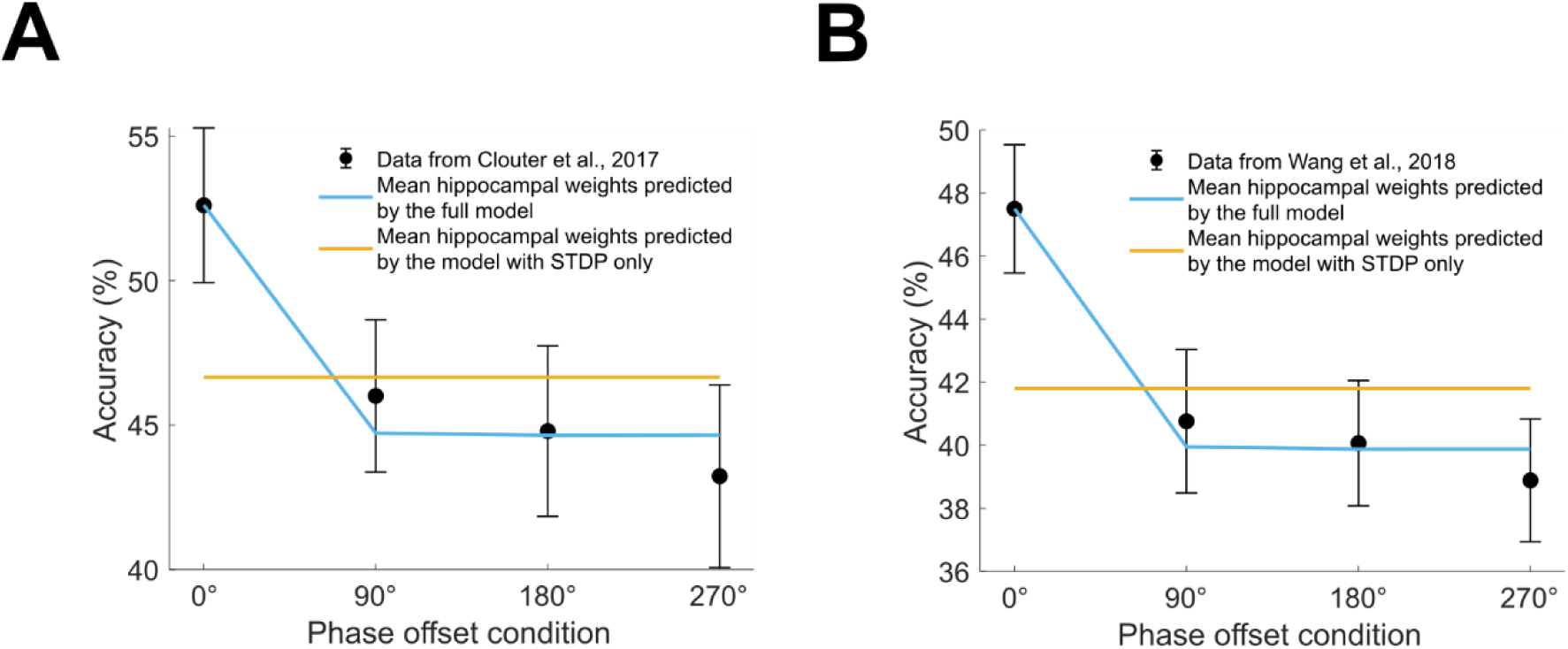
Model comparisons in recall performance between the full model and the STDP only model. ***A,*** Mean of weights from the Auditory to the Visual subgroup after learning simulated by two versions of model was fit to the data from Clouter et al. (2017). The full model was compared with an STDP only version of the model, which the range of input strength was between 0 and 1. ***B,*** Same as ***A***, but mean of weights from the Auditory to the Visual subgroup after learning simulated by two versions of model was fit to the data from Wang et al. (2018). All error bars represent SE.

## Notes

https://github.com/GP2789/Sync-deSync-model-TIME

### Competing Interest Statement

The authors have declared no competing interest.

